# Hamiltonian Monte Carlo with strict convergence criteria reduces run-to-run variability in forensic DNA mixture deconvolution

**DOI:** 10.1101/2022.02.15.480571

**Authors:** Mateusz Susik, Holger Schönborn, Ivo F. Sbalzarini

## Abstract

**Motivation:** Analysing mixed DNA profiles is a common task in forensic genetics. Due to the complexity of the data, such analysis is often performed using Markov Chain Monte Carlo (MCMC)-based genotyping algorithms. These trade off precision against execution time. When the default settings are used, as large as a 10-fold changes in inferred likelihood ratios (LR) are observed when the software is run twice on the same case. So far, this uncertainty has been attributed to the stochasticity of MCMC algorithms. Since LRs translate directly to strength of the evidence in a criminal trial, forensic laboratories desire LR with small run-to-run variability.

**Results:** We present a Hamiltonian Monte Carlo (HMC) algorithm that reduces run-to-run variability in forensic DNA mixture deconvolution by around an order of magnitude without increased runtime. We achieve this by enforcing strict convergence criteria. We show that the choice of convergence metric strongly influences precision. We validate our method by reproducing previously published results for benchmark DNA mixtures (MIX05, MIX13, and ProvedIt). We also present a complete software implementation of our algorithm that is able to leverage GPU acceleration, accelerating the inference process. In the benchmark mixtures, on consumer-grade hardware, the runtime is less than 7 minutes for 3 contributors, less than 35 minutes for 4 contributors, and less than an hour for 5 contributors with one known contributor.

## 1 Introduction

Investigators present at a crime scene identify and collect the available physical evidence. As a part of this evidence, DNA samples containing material from multiple contributors (i.e. *mixed DNA samples*) are often collected. The resulting short tandem repeat data is affected by stochastic events such as severe peak height imbalance, drop-outs, and drop-ins [3], especially in case of low-template samples. Manual analysis of the electropherograms (EPG) is unreliable and biased [39]. Therefore, laboratories rely on validated statistical software to solve the task of DNA mixture deconvolution [24].

The recommended metric [1] for reporting results of DNA mixture analysis is the likelihood ratio (LR):

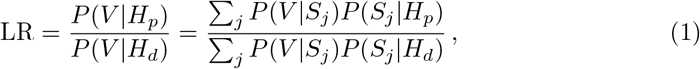

where *V* is the observed EPG, *S*_*j*_ represents a genotype set—a list of tuples denoting the allele designations of contributors. The summations are over all possible genotype sets *j. H*_*p*_ and *H*_*d*_ are the hypotheses of the prosecutor and the defendant respectively. A hypothesis assumes inclusion of certain contributors (suspect, victim, etc.) in the mixture, as well as the background allele frequencies of the populations the contributors allegedly belong to. Usually, the difference between the hypothesis of the prosecutor and the hypothesis of the defendant is the inclusion of the suspect in the former. In the setting we consider, the number of considered contributors is fixed beforehand. *P* (*S*_*j*_|*H*_*n*_) can be calculated based on the background frequencies of the alleles in the populations of interest for any hypothesis *H*_*n*_, *n* = {*p, d*} [8]. *Probabilistic genotyping* (PG) refers to the set of statistical methods used to compute LR for given EPG data.

In order to estimate *P* (*V* |*S*_*j*_), assumptions about the underlying data-generating process are made. These assumptions lead to a probabilistic model *P* (*V* |*M, S*_*j*_) with latent variables *M*. Models where probability is estimated based on the heights of the EPG peaks are called “fully continuous”. Two main approaches are used to infer such models: finding the most likely set of parameters by maximum likelihood estimation [7] or estimating the posterior [7, 33, 36]:

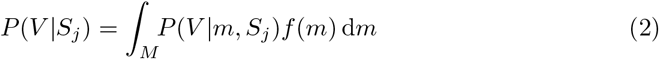

with the prior probability density function (PDF) of the parameters *f* (·). In an ideal scenario, LR is independent of any choices made by the laboratory technician and of any random confounding factors. In practice, however, LR depends on the variation in the samples from the crime scene, stochastic events occurring during DNA amplification, allele frequency sampling, parameter settings in the data-processing software^1^, hyper-parametrisation of the PG software, etc. [32]. Still, even when fixing all of these influences across identical runs on the same EPG data, residual *run-to-run variability* remains [5, 9, 13, 33, 36].

So far, this run-to-run variability has been attributed to the inherent stochasticity of the Markov Chain Monte Carlo (MCMC) methods used to estimate mixture model parameters [9, 11]. However, as we show here, the apparent run-to-run variability is more likely caused by the choice of convergence criteria used in the MCMC sampler. This is supported by our results demonstrating that run-to-run variability can be reduced when using an MCMC method with strict convergence criteria. For this, we formulate a probabilistic model of DNA mixture deconvolution that only has continuous degrees of freedom, marginalising over the discrete dimensions. While such marginalised models can be properly convergence controlled, they are generally more expensive to solve. As we show here, though, the intrinsic structure in the problem can be exploited by Hamiltonian Monte Carlo (HMC), maintaining the runtimes of conventional MCMC solutions. We show that the strict convergence criteria afforded by our method significantly reduce run-to-run variability. We further present data structures that efficiently handle the combinatorial growth in the number of genotype sets with increasing numbers of contributors, and we present a GPU-enabled implementation of DNA mixture deconvolution.

### 1.1 Precision of DNA mixture deconvolution

We define *precision* of DNA mixture deconvolution as the inverse variance between results of runs with identical hyper-parametrisation on the same EPG data for the same hypotheses. Precision has to be considered in addition to the accuracy of a PG system [32], as also the authors of PG algorithms note [27]:

> “The argument is that the existence of variability *[across PG runs — our note]* raises doubts about whether any of the results should be accepted.” [27]

Courts are often unaware of run-to-run variability, as expert witnesses usually report a single LR number [25]. The issue is even more severe when the verbal scale for reporting LRs is used [2, 23]. European Network of Forensic Science Institutes [23] suggest a scale that defines LR between 100 and 1000 as “moderately strong support” and LR between 1000 and 10 000 as “strong support”. Let us assume that we use a PG system, which, for the given case, outputs results from a normal distribution: log_10_ LR ∼ *𝒩* (2.3, 0.5). A single run of the software would give “strong support” in ≈73% of cases. A technique that reports confidence intervals [8, 10], however, would provide the conservative answer of “moderately strong support”. This highlights the importance of high precision (i.e. low variance) in PG results.

The precision of available commercial solutions has been quantified in several studies [5, 9, 13, 33, 36]. A standard deviation of LR of *>* 10^4^ has been reported between identical runs on a three-contributor mixture (sample ‘3-2’) when the TrueAllele® software was used [5]. Results obtained with the STRmix™ software displayed 10-fold LR difference across runs [9, 13]. In order to increase precision, it has been recommended to increase the number of MCMC iterations, at the expense of a larger computational runtime [13, 34].

To determine when to terminate an MCMC sampler in Bayesian inference, convergence criteria are used [26]. The most popular criterion is the univariate *Gelman-Rubin* (GR) diagnostic [14, 26], which compares pooled and within-chain variances of samples to indicate possible convergence. For given model parameters, this diagnostic has a value close to 1 if the samples from different chains result in similar estimates for the marginal distribution. Since actual convergence can not be quantified as long as the true posterior distribution is not known, convergence criteria measure the stability of samples, and the term “convergence” in MCMC is always relative to the chosen test statistic.

Some of the available PG software solutions provide users with convergence diagnostics. STRmix™ [36] for example calculates the ratio of pooled and within-chain variances of the likelihood of the model (personal communication, Kevin Cheng, Institute of Environmental Science and Research, Ltd., Wellington). GenoProof Mixture [28] reports the univariate GRs for the continuous parameters. By default, both software solutions use a predefined constant number of post burn-in samples and then report the value of the diagnostic to the user. If the desired threshold (by default in STRmix™, 1.05 in GenoProof Mixture) of the convergence diagnostic has not been achieved, the software offers an option to run additional iterations. The default GR threshold should be rather low. The authors of the diagnostic state [26]:

> “The condition of GR near 1 depends on the problem at hand; for most examples, values below 1.1 are acceptable, but for a final analysis in a critical problem, a higher level of precision may be required.” [26]

Providing evidence in court should be considered a critical problem, as the consequences of wrong or doubtful answers are significant [30]. Other scientists researching convergence diagnostics therefore noted [40]:

> “We argue that a cutoff of GR ≤1.1 is much too high to yield reasonable estimates of target quantities.” [40]

This seems even more important since the statistical models used in both of these software tools combine continuous (e.g. peak intensities) and discrete (e.g. genotype sets) dimensions. However, GR can not monitor convergence in discrete dimensions. It is therefore possible that convergence is deduced purely from the continuous parameters, while the genotype set distributions may not have converged at all, offering a possible explanation for the large run-to-run variability observed despite low GR thresholds. In our model, we avoid the issue of assessing convergence of genotype sets by marginalising them out.

### 1.2 Trade-off with execution time

In forensic DNA mixture deconvolution, computational runtime is of great importance, since:

- Laboratories might have to run software multiple times with different hyper-parametrisations in order to check the robustness of the results or to test hypotheses with different numbers of contributors, different analytical thresholds, different priors, etc.
- Laboratories might want to quantify the precision of the results over several identical replicates.
- Forensic laboratories are often working under time pressure, e.g., if cases attract great media attention or laws limit detention time without charges.

In addition to the efficiency of the software implementation, there are multiple factors that influence the execution time, including the number of contributors, the number of alleles per locus, the techniques used to limit the number of considered genotype sets, the convergence criteria, the choice of the optimisation problem (maximum likelihood vs. Bayesian inference), the specification of the model, the priors used, etc.

In general, there is a trade-off between the precision and runtime. Lower runtimes can trivially be achieved by running a smaller number of MCMC iterations, at the expense of precision. Achieving perfect precision is theoretically possible, if runtime is unbounded, by integrating over the latent variables (see Eq. 2).

The model presented here integrates over the discrete dimensions, i.e., marginalises over genotype sets in order to be able to properly monitor convergence. In a standard MCMC sampler, this would lead to greatly increased runtimes, hampering practical applicability. As described below, however, it turns out that the structure of the resulting search space permits efficient exploration by HMC. Thanks to the resulting increase in sample efficiency, we are able to use a strict GR threshold of 1.05 with similar or faster runtime than existing solutions.

## 2 Materials and methods

We base our probabilistic genotyping model on the work by Taylor et al. [36], due to the large number of studies that describe and evaluate this model (e.g. [7, 17, 20, 36, 38]). The main assumption in this model is the log-normal distribution of the ratio of the observed EPG peak heights to the peak heights predicted by the generative model (called “expected” peak heights). The generative model consists of several steps, as illustrated in Fig. 1: First, the expected contributions are computed for each genotype set from the considered set of parameters. Then, peak stutter models are applied to predict expected peaks. The next step is to calculate the standard deviation 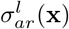 of the log-normal distribution. Finally, the likelihood of the observed data given the parameters is calculated as the product of the likelihoods of all peaks. The model handles stochastic dropout and drop-in events. To provide a mathematical formulation of the model, we denote:

**Figure 1.**
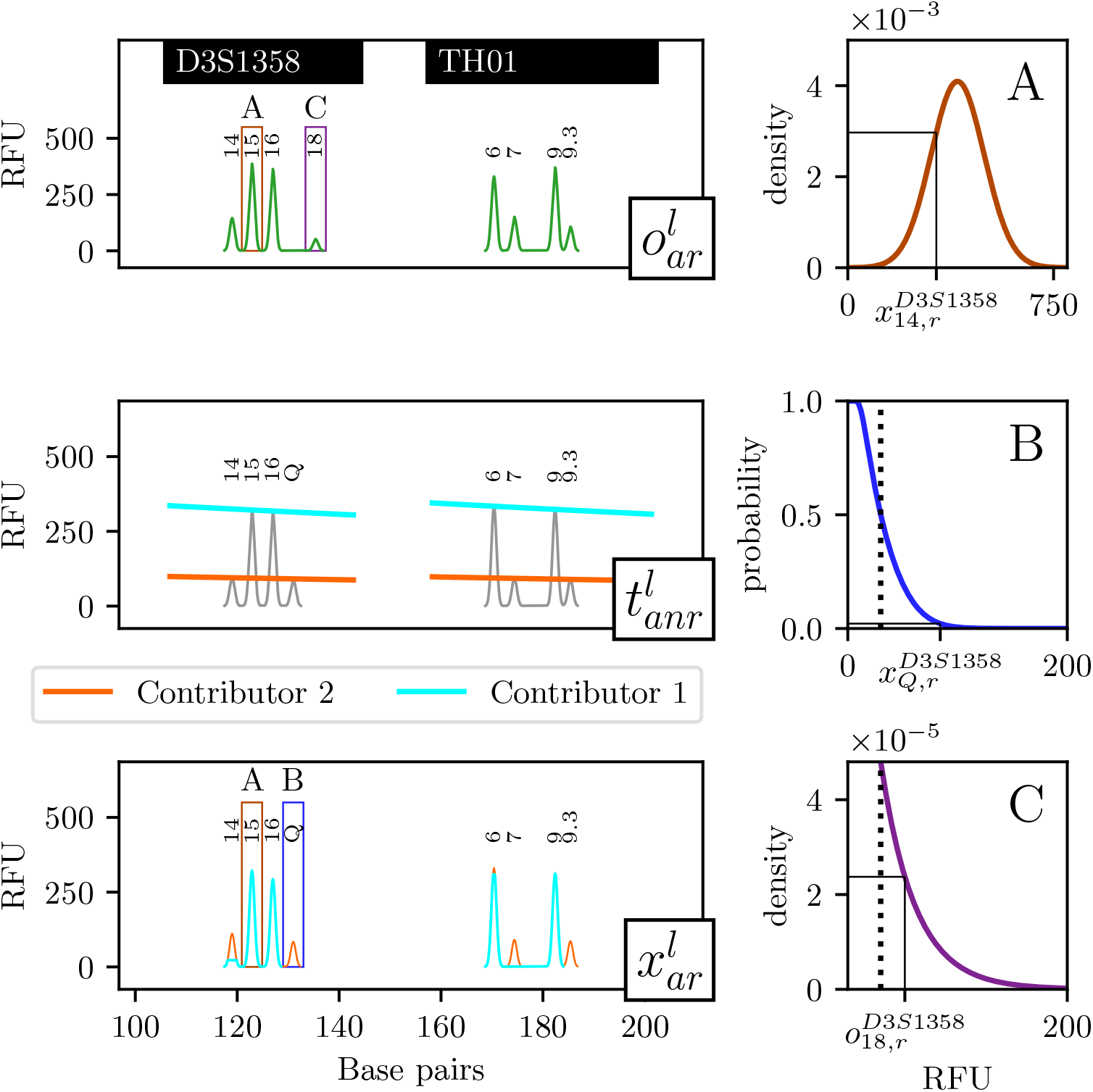
Illustration of the present probabilistic genotyping model. Top left: the observed peaks 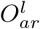 at two selected loci from the green channel of the analysed mixture. Centre left: We analyse the genotype set [(15,16),(14,Q)] for locus D3S1358 and [(6,9),(7,9,3)] for locus TH01. The catch-all dropout allele *Q* denotes any dropped-out peaks. The lines show the expected contributions for all alleles 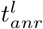. The function is decreasing within a locus due to the effect of decay modelling. TH01 locus was modelled with a larger amplification efficiency parameter than D3S1358. Bottom left: Expected peaks 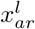 are created by applying stutter ratios to 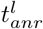 for two contributors. In this example, we consider only backward stutter for illustration purposes. Composed peaks (i.e. those that consist of both allelic and stutter contributions) are 14 and 15 at D3S1358, and 6 at TH01. Top right: PDF of the log-normal model for peak 15 at D3S1358 as a function of the expected peak height. Centre right: dropout probability as a function of expected peak height. Bottom right: drop-in probability as a function of observed peak height. Peak 18 at D3S1358 is a drop-in, since it is observed but not expected in the considered genotype set. The dotted lines denote the analytical thresholds.

- 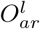: random variable of observed peak height at locus *l*, allele *a*, replicate *r*;
- 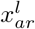 : expected peak height at locus *l*, allele *a*, replicate *r*;
- *f*_*X*_ : the PDF of a random variable *X*;
- *Q*: the “catch-all” dropout allele [36];
- 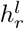 : the analytical threshold of the EPG for locus *l* and replicate *r*;
- N (0, *s*^2^): a normal distribution with mean 0 and standard deviation *s*;
- (*M, N*): alleles of a contributor at a single locus, e.g (12,13).

We consider the posterior probability

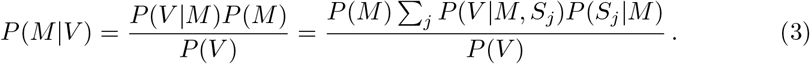

As this analysis is independent of the hypothesis, we do not favour any genotype and treat them as equally likely. In Bayesian inference, evidence is usually neglected as it is too expensive to compute and constant w.r.t. model parameters. We thus obtain the unnormalised posterior

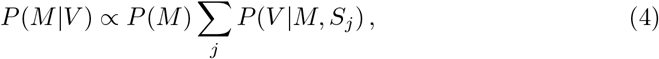

which we use for estimating *P* (*V* |*S*_*j*_) (see Eq. 2). We assume peak heights to be conditionally independent given *S*_*j*_ and *M*, and alleles of a contributor in different loci to be independent from each other. The resulting model is a function 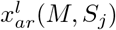 of *M* and *S*_*j*_. The likelihood of the observed EPG given parameters *M* and a genotype set *S*_*j*_ is then:

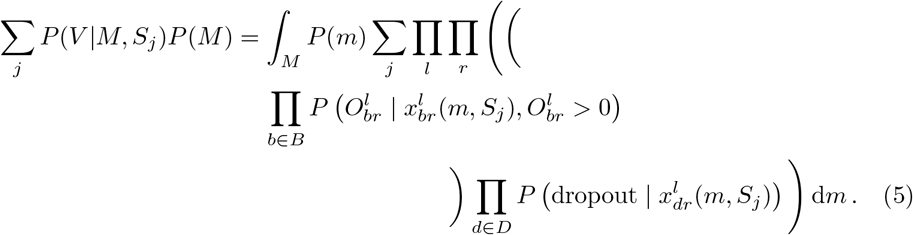

The inner multiplications are performed over the set *B* of observed peaks and the set *D* of hypothetical dropout peaks. This model formulates separately the relative likelihood of observed peaks and the probabilities of dropout events. In the following, we abbreviate the notation for 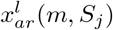 to 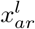.

### 2.1 Observed peaks

In case a peak is observed (i.e.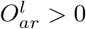) our model considers a mixture distribution. The mixture combines sub-models for peaks that are expected (*f*_*Z*_) and for drop-in events 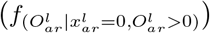 :

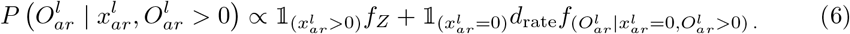

For the drop-in events, we use the model introduced by Euroformix [7]:

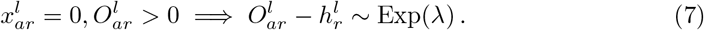

Two hyper-parameters based on the level of noise in negative controls are required: the drop-in rate *d*_rate_ and the *λ* of the exponential distribution. For the expected peaks, we assume a log-normal distribution following previous works [36]:

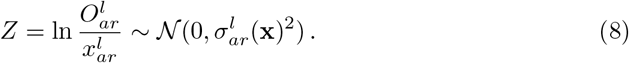

The standard deviation of this distribution depends on the components of the expected peaks and their heights:

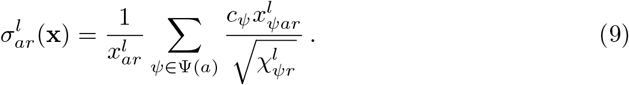

We define *ψ* ∈ Ψ(*a*) = {*a* + 2, *a* + 1, *a, a* −1}. This means that for a single allele *a*, we consider contributions from the allelic peak *a*, the backward stutter from the *a* + 1 peak, the forward stutter from the *a* − 1 peak, and the double backward stutter from the *a* + 2 peak. We use one parameter for allelic peak standard deviation (*c*_*ψ*_ = *c*_*p*_ when *ψ* = *a*) and a different one for stutter peak standard deviation (*c*_*ψ*_ = *c*_*s*_ when *ψ* ≠ *a*). Additional parameters for different types of stutter could be introduced without significantly changing the model. The expected peak heights then are:

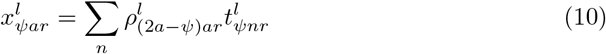

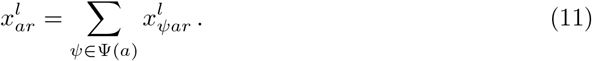

The sum in Eq. 10 is over the assumed contributors. Finally, 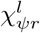 models the fact that peak variance is inversely proportional to peak height [15]. The rationale behind the formula is explained in Chapter 1 of the Supplementary Material:

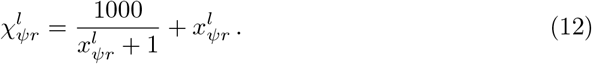

Equation 10 includes the normalised stutter ratios *ρ* and the product contributions from a contributor 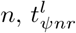. To obtain *ρ*, the unnormalised stutter ratios *π* are deduced from unambiguous profiles. They are modeled with linear regressions based on allele designation or longest uninterrupted sequences [36]. Normalization is subsequently required, since multiple types of stutter are considered at the same time:

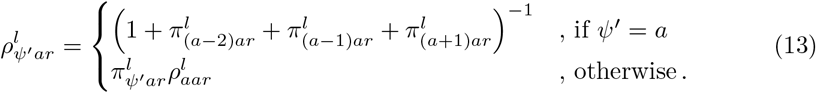

The product contribution at a selected allele *a* is defined as:

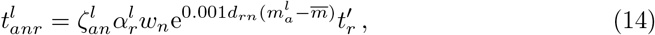

where 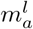 is the molecular weight of the allele, and 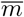 is the average of the largest and smallest observed molecular weights on the EPG within the called peaks. The integer 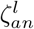 is 1 if the genotype of contributor *n* includes allele *a* in locus *l*, and 0 otherwise. It is equal to 2 in case of a homozygote. The scalars 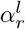 are the locus-specific amplification efficiencies (LSAE), *w*_*n*_ are the weights of the contributors that sum up to 1, *d*_*rn*_ are the decay parameters, and 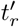 is the total allelic product expressed in relative fluorescent units (RFU).

### 2.2 Dropout events

In case of a dropout, the peak is unobserved because it is below the analytical threshold. This corresponds directly to

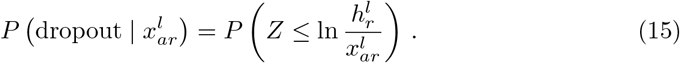

See Supplementary Material section 1 for details.

### 2.3 Parameters of the model and priors

We define the prior probability of the parameters as:

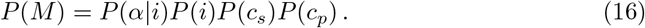

Here, *P* (*α*|*i*) is the prior ln 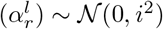 that prevents the amplification efficiencies from drifting away from 1 too far. The prior variance *i*^2^ ∼ Exp(*σ*_*α*_), where *σ*_*α*_ is a hyper-parameter to be optimised by the laboratory [12]. *P* (*c*_*s*_) and *P* (*c*_*p*_) are optional priors on the peak height standard deviation parameters, which are also present in STRmix™ [12].

The free parameters to be explored by the sampling algorithm then are: 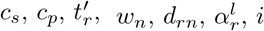. We reduce the number of parameters if multiple replicates are performed using the same kit. In such a case, the analysis shares the LSAEs 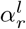 across the replicates. Let us denote by *l*_all_ the resulting total number of LSAE parameters. Then, there are 2 + |*r*|+|*n*|+ |*r · n*|+*l*_all_ parameters overall, whose values have to be estimated (one of the weights is trivial when all others are set).

### 2.4 Other considerations

In order to provide conservative estimates of LR, we use Balding-Nichols sub-population correction [4, 8], and we report sub-source LRs [37] unless specified otherwise. We consider dropout allelic contributions as separate peaks, i.e., (Q,Q) is considered heterozygous. The total number of genotype sets is reduced by considering at most one drop-in per locus and using a drop-in cap. Peaks which are in stutter position and are not included in the genotype set definition are considered drop-ins if they are abnormally tall w.r.t. the origin, see the maximum stutter ratios in Table 2.

### 2.5 Sampling algorithm

The marginalised (over genotype sets *S*_*j*_) PG model from Eq. 4 is prohibitively costly to solve with MCMC, due to the large number of log-probabilities that need to be calculated when all possible genotype sets are considered. We therefore use an adaptive-proposal sampler that has been successfully used in other fields: Hamiltonian Monte Carlo (HMC) [22]. The difference between HMC and MCMC is how the proposal distribution is chosen. HMC simulates physical (i.e. Hamiltonian) system dynamics instead of choosing a random point from the neighbourhood of the current sample. This renders HMC very efficient for posteriors with multi-modal or multi-funnel shapes, significant parameter correlations, and/or high dimensionality [6]. Unlike STRmix™ and GenoProof Mixture, which only change the value of a single continuous parameter in each iteration, our sampler considers multi-variate moves. The proposal distribution for these moves is dynamically adapted across iterations. It is determined in each iteration based on the local gradient of the log-probability of the model [22]. This is possible because our model is differentiable, as the discrete genotype sets are marginalised out.

To compute the model gradients, we rely on the proven automatic differentiation framework in TensorFlow Probability [21], where HMC is also implemented. Thanks to the portability of the TensorFlow library, we can provide both CPU and GPU versions of our algorithm. However, a naïve TensorFlow implementation would perform below expectations due to the combinatorial growth of the number of possible genotype sets with increasing number of contributors. We therefore introduce an important performance improvement: a deduplication system that can be used with any model that considers all genotypes in a single iteration (e.g. our work, Euroformix). Additional performance optimisations are described in Chapter 3 of the Supplementary Material.

For the deduplication system, we consider a single locus *l*. The expected peak heights 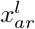 and the standard deviation 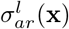 of their distribution (Eq. 8) depend on the genotype sets. If we consider the output for a single allelic position, these values depend only on the continuous parameters as well as the counts of alleles of individual contributors from the selected position and the stutter positions. When multiple genotype sets are considered, the same values are needed multiple times. As an example, consider the locus shown in Fig. 2 and the likely genotype sets from Table 1 for two contributors. Computing every expected peak (and the likelihoods of observing the ratios between the observed and expected peaks) for genotype set {(10,13), (12,15)} entails computations that are also identically required for the other genotype sets. Our deduplication system ensures that each such computation is performed only once, and the result is cached and reused. This leads to large savings in multi-contributor mixtures. As an example, deduplication reduced the number of peak predictions in one locus of a 4-contributor mixture from 44 473 to 524. We use deduplication during both the calculation of the log-probability and the calculation of the HMC gradient.

**Table 1.** Three likely genotype sets *S*_*j*_ and the resulting contributions to the peaks of the locus in Fig. 2 from two contributors. *C*_*n,a*_ = *ρ*_*aa*1_*t*_*an*1_ is the contribution of contributor *n* from a single copy (i.e. *r* = 1) of allele *a*. 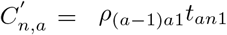 is the contribution from stutter originating from allele *a*. For simplicity, only single-backward stutter and a single replicate are considered. Duplicate entries are highlighted with the same colour.

| $S_j$ | 10 | 12 | 13 | 14 | 15 | Q |
| --- | --- | --- | --- | --- | --- | --- |
| (15,Q), (10,13) | $C_{2,10}$ | $C'_{2,13}$ | $C_{2,13}$ | $C'_{1,15}$ | $C_{1,15}$ | $C_{1,Q}$ |
| (10,Q), (13,15) | $C_{1,10}$ | $C'_{2,13}$ | $C_{2,13}$ | $C'_{2,15}$ | $C_{2,15}$ | $C_{1,Q}$ |
| (10,13), (12,15) | $C_{1,10}$ | $C'_{1,13} + C_{2,12}$ | $C_{1,13}$ | $C'_{2,15}$ | $C_{2,15}$ | 0 |

**Figure 2.**
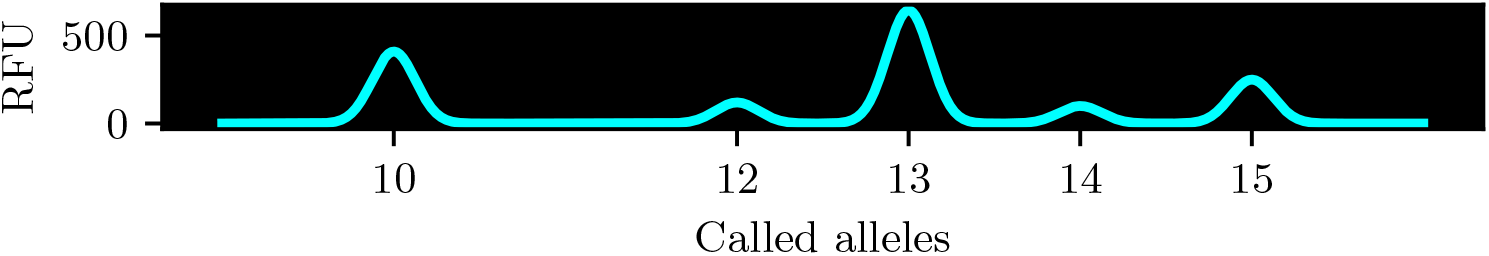
Example of a locus from a DNA mixture with two contributors.

The deduplication system works as follows: Before a run is performed, we precompute the indices of all duplicate entries in the functions and create two data structures, one containing the information required to evaluate the deduplicated expected peak heights, and one containing the indices for a gather operation to be performed after the log-probabilities of the deduplicated peaks have been calculated. This gather operation then unfolds the deduplicated results into a matrix that stores the log-probabilities per replicate, allelic position, chain, and genotype set.

## 3 Results

Reliable forensic genotyping should be free of false inclusion and false exclusion. Moreover, it should be precise and have low inference runtime. We quantify these aspects for our proposed solution on publicly available test mixtures from previously published benchmarks: the ProvedIt dataset [34] and the MIX05 and MIX13 studies [18]. For the GlobalFiler™ mixtures from the ProvedIt dataset, we use the hyper-parametrisation suggested by Riman et al. [34]. For the MIX05 and MIX13 datasets, we use the hyper-parametrisation from Buckleton et al. [16]. For MIX05, MIX13, and precision studies we use the *FBI extended caucasian population* [31] genetic background model. For the challenging mixtures of Subsection 3.3, we use the *NIST 1036-Caucasian* background allele frequencies [35].

We denote the contributors in the hypotheses by plus-delimited strings. U stands for an unknown contributor, W is a witness, V is a victim, and all other entries denote suspects. The hypothesis V+W+S+U for example has 4 contributors: the victim, the witness, the suspect, and one unknown person. In all benchmark cases, the defendant’s hypothesis is the prosecutor’s hypothesis with the suspect replaced by an unknown contributor. All benchmark mixtures were created in laboratories with known ground-truth genotypes of the contributors.

The linear stutter models are fit on single-source profiles (forward, backward, and double-backward stutter for ProvedIt and MIX13) or on data provided by the kit manufacturer (only backward stutter for MIX05). The stutter models are available in Chapter 4 of the Supplementary Material.

All experiments are performed on affordable hardware in the cloud. We use NC8as T4 v3 instances from Azure Cloud (8 vCPUs, Nvidia Tesla T4 GPU, 16 GB RAM). An exception has been made for ProvedIt Sample 3, which does not fit within 16 GB of RAM. For this case we rented a Google Cloud virtual machine with a Nvidia A100 GPU.

### 3.1 Accuracy: MIX05 and MIX13 benchmarks

We first benchmark the performance of our method on inter-laboratory studies organised by NIST: MIX05 and MIX13 [18]. For MIX05, we analyse simultaneously replicates from different kits: ABI’s COFiler, ABI’s SGM Plus, Promega’s Powerplex 16, and ABI’s Profiler Plus. For MIX13, we follow the published studies in using only ABI’s AmpFLSTR IdentiFiler Plus replicate. All cases are analysed with a global analytical threshold of 50 and the ground truth number of contributors with two exceptions: Case 5 from MIX13 is also analysed with 3 contributors (since most laboratories taking part in the original study estimated this number), and Case 2 from MIX13 uses an analytical threshold of 30 (following the recommendation from NIST). For the capillary electrophoresis fragment analysis files, we use default GeneMapper™ ID-X 1.4 analysis settings. The results are presented in Table 3. Our algorithm reproduces most of the results of other solutions, suggesting its validity. Similar to other solutions, our algorithm provides more conservative LR values when a smaller number of contributors is chosen [16]. The only case in which our model provided LR larger than 1 for a false suspect is S05C in MIX13 Case 5. The genotype of this suspect had been deliberately constructed to share alleles with the true contributors in every locus. 74 out of 108 laboratories have included this suspect in the original study [18]; our method excludes it in a 3-contributor scenario. For MIX13 Case 4, our algorithm provides a higher LR than the reciprocal of the random match probability. An explanation for this behaviour is given in Subsection 2.1 of the Supplementary Material.

**Table 2.**
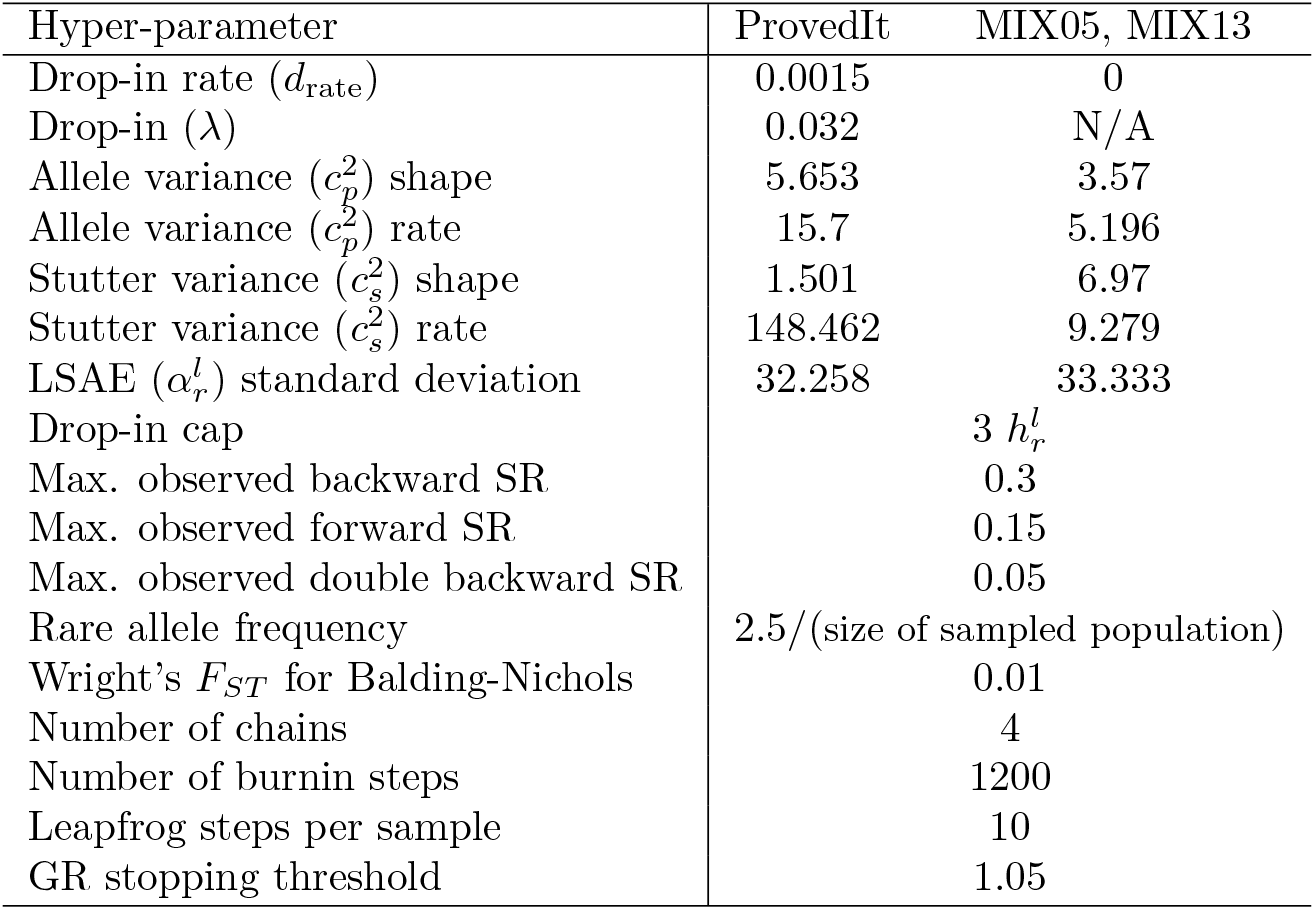
Hyper-parameter values used in the present benchmarks. SR stands for stutter ratio. The values of the variance rates and the standard deviation of the locus-specific amplification efficiency (LSAE) were adjusted to our formulation of the model (i.e., natural logarithm instead of log_10_, 1/mean for LSAE standard deviation).

**Table 3.** Accuracy of our method. Average LRs over 10 runs for our method (“HMC”) in comparison with other solutions on the MIX05 and MIX13 benchmarks with known ground truth. In most cases, all tested methods correctly return LRs larger than 1 for ground-truth inclusion or smaller than 1 for ground-truth exclusion. For MIX13, we compare with the reported results of Euroformix (“EFM”) version 1.11.4 and STRmix™ [16]. The TPOX locus was ignored in the MIX05 Case 4, due to a tri-allelic pattern of the perpetrator.

| Case | $H_p$ | Truth | $\log_{10}$ LR | | |
| --- | --- | --- | --- | --- | --- |
|  |  |  | HMC | EFM | STRmix™ |
| MIX05 Case 1 | Perpetrator+U | Incl. | 19.34 | - | - |
| MIX05 Case 2 | Perpetrator+U | Incl. | 25.67 | - | - |
| MIX05 Case 3 | Perpetrator+U | Incl. | 22.32 | - | - |
| MIX05 Case 4 | Perpetrator+U | Incl. | 10.44 | - | - |
| MIX13 Case 1 | V+S01A | Incl. | 20.15 | 20.18 | 20.15 |
| MIX13 Case 2 | S02A+U+U | Incl. | 17.03 | 17.28 | 16.98 |
| MIX13 Case 2 | S02B+U+U | Incl. | 7.50 | 7.88 | 7.26 |
| MIX13 Case 2 | S02C+U+U | Incl. | 5.41 | 6.11 | 5.83 |
| MIX13 Case 2 | S02D+U+U | Excl. | -16.18 | -2.36 | -14.03 |
| MIX13 Case 3 | V+W+S03A | Incl. | 7.87 | 6.82 | 7.69 |
| MIX13 Case 3 | V+W+S03B | Excl. | $-\infty$ | $-\infty$ | $-\infty$ |
| MIX13 Case 4 | V+S | Incl. | 20.23 | 19.91 | 20.15 |
| MIX13 Case 5 | S05A+U+U | Incl. | 3.38 | 9.26 | 3.45 |
| MIX13 Case 5 | S05B+U+U | Incl. | 1.61 | 9.38 | 3.32 |
| MIX13 Case 5 | S05C+U+U | Excl. | -8.66 | 6.45 | -9.22 |
| MIX13 Case 5 | S05A+U+U+U | Incl. | 6.20 | - | - |
| MIX13 Case 5 | S05B+U+U+U | Incl. | 5.96 | - | - |
| MIX13 Case 5 | S05C+U+U+U | Excl. | 2.63 | - | - |

### 3.2 Precision: ProvedIt benchmark

Next we analyse the precision of our method in comparison with the state of the art on the ProvedIt inter-laboratory benchmark [9]. We use the same analytical thresholds as Kelly et al. [29] and report sub-sub-source LRs following Ref. [9].

We focus on ‘Sample 1’ cases, which were previously used to determine the precision of STRmix™ [9].^2^ Four different analysis methods of GeneMapper™ are used in comparison (called A,B,C, and D). The results are shown in Fig. 3. For STRmix™, a 10-fold run-to-run variability in the LRs is observed with the default stopping criterion (blue), which has been attributed to the stochastic nature of MCMC [9]. In our HMC method, we check the GR diagnostic every 300 iterations and stop when it is below 1.05 for all parameters. This results in roughly 10 orders of magnitude reduction in the LR standard deviation (cyan). For GeneMapper™ analysis method A, the standard deviation of log_10_ LR is 10.08 times lower in our method than in STRmix™ [9] (Fig. 3A). For analysis method B (Fig. 3B), it is 8.76 times lower, for analysis method C 10.99 times lower (Fig. 3C), and for analysis method D 10.76 times lower (Fig. 3D).

**Figure 3.**
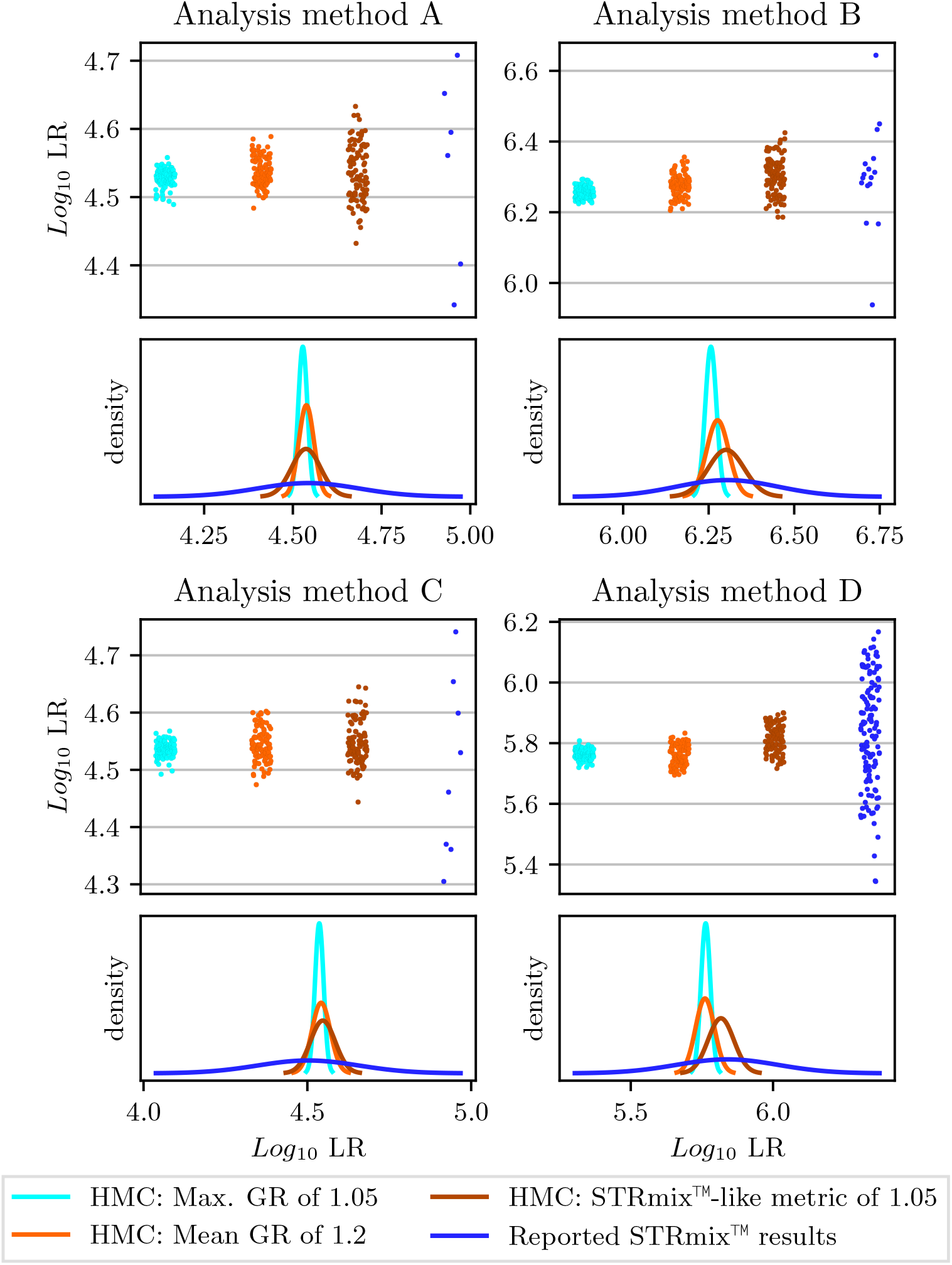
Precision of our HMC method using different stopping criteria in comparison with STRmix™. We use four different GeneMapper™ analysis methods (A, B, C, D). Every case is run 100 times, and the resulting per-run LRs are shown as dots. The corresponding maximum likelihood estimations of a normal distribution are shown in the plots below. With standard stopping criteria, our method (cyan) reduces the standard deviation of log_10_ LR around 10 fold over STRmix™ (blue). From the published STRmix™ results, we ignored the result provided by participant L4A1 (no known contributor included) and the second run of participant L1A1 (missing in the plots of the original work [9]).

In the same Fig. 3, we also quantify the influence of the convergence diagnostic on the precision of our algorithm by testing two alternative stopping criteria: In the first, we calculate the mean of GR values and stop when this mean is *<*1.2 (orange). In the second, we use the same convergence metric as STRmix™, but with our threshold value of 1.05 (brown). Due to the efficiency of our HMC sampler, we were unable to simulate chains that resulted in values of this metric approaching 1.2. Interestingly, however, we find that the STRmix™ criterion was sometimes satisfied with threshold 1.05 when some GR values were still *>*1.2.

Taken together, these results show that our approach is able to significantly improve precision over the STRmix™ method, and that this improvement is due to the stricter convergence criteria, as enabled by our purely continuous model with discrete dimensions marginalised out.

### 3.3 Performance: computationally challenging mixtures

Finally, we quantify the performance of our HMC method on the most challenging cases of mixtures with 3 to 5 contributors. This serves to test our approach on cases that are not simple to resolve and that are challenging both from a precision point of view and for computational runtime. For these cases, we use the set of low analytical thresholds from Riman et al. [34].

We analyse 4 mixtures: Sample 1 and Sample 2 from Bright et al. [9] analysed here without a known contributor (referred here to as Sample 1b and Sample 2b, respectively), and two following 5-person mixtures from ProvedIt with one known contributor:

- A05_RD14-0003-30_31_32_33_34-1;1;1;1;1-M3I22-0.315GF-Q1.3_01.15sec (named here Sample 3, known contributor is Contributor 31)
- E04_RD14-0003-48_49_50_29_30-1;1;2;4;1-M2d-0.279GF-Q2.1_05.15sec (named here Sample 4, known contributor is Contributor 29)

For each case, we construct all possible prosecutor hypotheses with 1 suspect. We also construct the same number of false hypotheses by choosing the suspects randomly from the NIST 1036 U.S. Population Dataset [35]. To quantify precision, we run our method 10 times for each ProvedIt case. The results are shown in Fig. 4. In all cases, our algorithm correctly classifies contributors and non-contributors with high precision. In all but one case where the true contributors are considered, the difference between extremal values of log_10_ LR is under 0.2.

**Figure 4.**
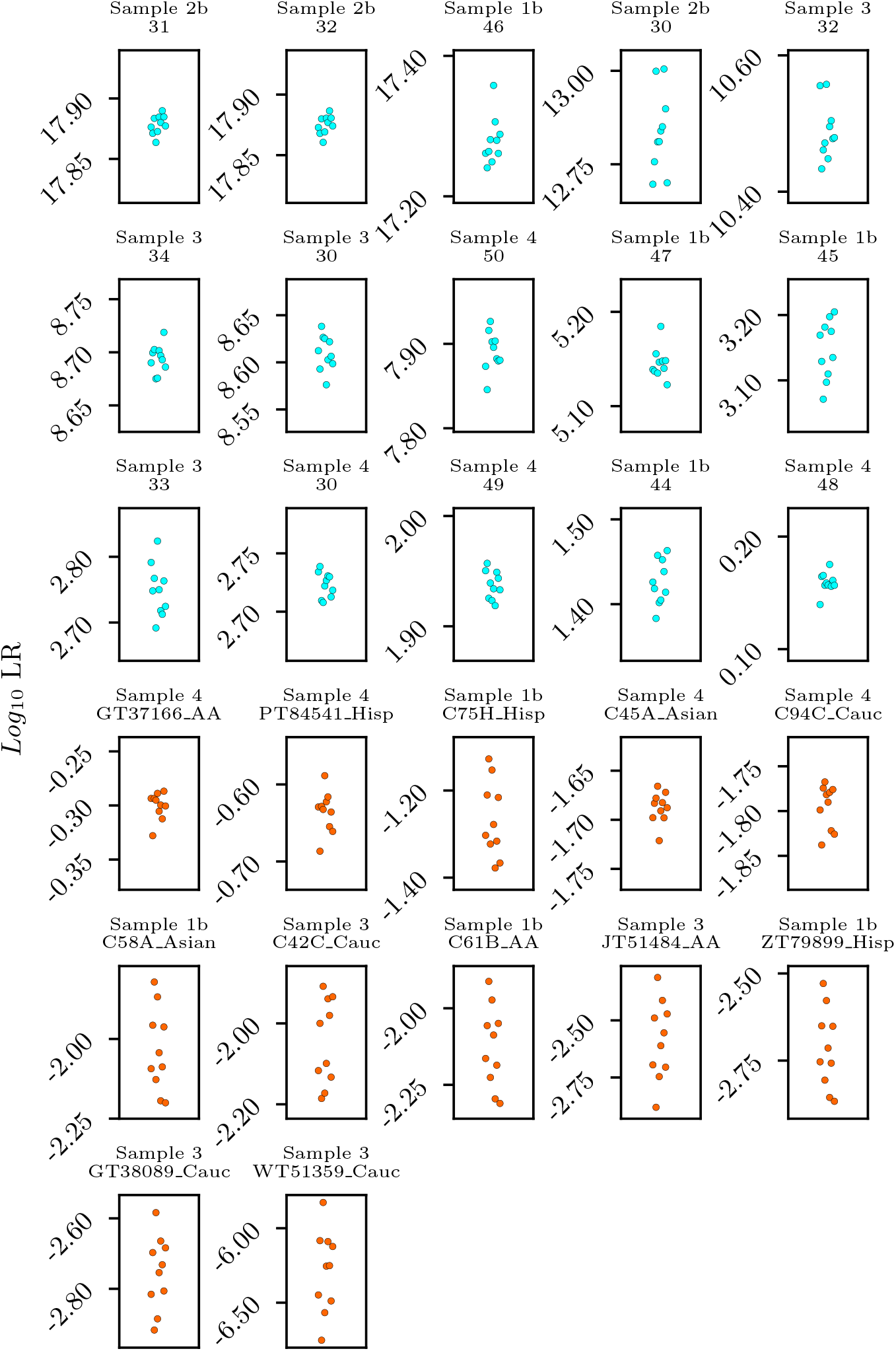
Precision for computationally challenging ProvedIt mixtures. Panel titles indicate the case (top row) and the suspect (bottom row). Each combination is run 10 times, and we plot sub-source log_10_ LR. In all but one case where the true contributors is considered the difference between the highest and the lowest log_10_ LR is less than 0.2. One of the exceptions is Sample 2b when the suspect is Contributor 30. This is further analysed in Chapter 2 of the Supplementary Material. We do not plot exclusion scenarios with LR = 0 for all runs. These are: Sample 2b – non-contributors C18C Cauc, C99B AA, and ZT80925 Hisp; Sample 3 – non-contributors GT38089 Cauc, JT51484 AA, and WT51359 Cauc. Inclusion cases with correct suspect are plotted in cyan, correct exclusion cases in orange.

The inference runtimes on the benchmark cloud instances are shown in Table 4. We show the results for all the mixtures we analysed with 3 or more unknown contributors. The results are better than the reported runtimes of previous versions of PG software solutions [13]. This suggests that despite the increased computational complexity of our marginalised model, the efficiency of HMC sampling and the efficient GPU implementation recover state-of-the-art runtimes as required for practical use of the method.

**Table 4.** Inference runtimes. We report the average (over 10 repetitions) inference times on a single-GPU cloud instance for the listed cases with different numbers of contributors (NoC) in minutes (m) and seconds (s).

| Case | NoC | NoC known | Inference time |
| --- | --- | --- | --- |
| MIX13 Case 2 | 3 | 0 | 3m 59s |
| MIX13 Case 3 | 3 | 0 | 1m 59s |
| MIX13 Case 5 | 3 | 0 | 4m 37s |
| MIX13 Case 5 | 4 | 0 | 31m 15s |
| ProvedIt Sample 1 | 4 | 1 | 4m 13s |
| ProvedIt Sample 1b | 4 | 0 | 19m 39s |
| ProvedIt Sample 2b | 3 | 0 | 6m 55s |
| ProvedIt Sample 3 | 5 | 1 | 59m 55s |
| ProvedIt Sample 4 | 5 | 1 | 27m 15s |

## 4 Conclusions and future work

High precision, i.e. low run-to-run variability, of the results provided by probabilistic genotyping methods is key to building trust and to ensuring reliable discriminatory power of the analyses. While run-to-run variability has previously been attributed to the inherent stochasticity of MCMC algorithms [9, 11], we have shown that it can be significantly reduced by an adjusted model formulation and stricter convergence criteria. We hypothesised that the convergence of probabilistic genotyping models is hard to assess if they contain both continuous (e.g. peak intensities) and discrete (e.g. genotype sets) dimensions. We therefore presented a model where the discrete dimensions are marginalised out, leading to a purely continuous and differentiable formulation. Thanks to the differentiability of our model, we were able to use Hamiltonian Monte Carlo (HMC) to achieve state-of-the-art inference runtimes with GPU acceleration.

The benchmark experiments presented have shown a reduction in the standard deviation of the resulting log-likelihood ratios by around an order of magnitude when using our method compared to the state-of-the-art STRmix™ software. They have also provided validation of the inference results against known ground truth by close reproduction of previously published results.

In the future, we plan to compare our method with other algorithms on the ProvedIt benchmarks (e.g. [19, 34]), provide a comparative analysis of the two main algorithmic approaches (Bayesian inference vs. maximum likelihood estimation) when the same probabilistic model is used, and work on further improvements of the model.

In addition to the run-to-run variability of the inference algorithm, the overall precision observed on a sample in the laboratory also depends on multiple other factors, including the frequency of the suspect’s genotype in the background population, the proportion of the suspect’s template, the quality of the sample, and the hyper-parametrisation of the data-processing methods. All of these must therefore be fixed when comparing different probabilistic genotyping algorithms. However, it might be insightful to explore which of these factors have the largest influence on the precision of final results, and to bound the precision in the worst case.

## Acknowledgements

We thank Dr. Sachin Krishnan (Center for Advanced Systems Understanding, Gorlitz) and Zofia Dziedzic (University of Wroclaw) for discussions on improving the mathematical notation, and Kevin Cheng (Institute of Environmental Science and Research, Ltd., Wellington) for valuable insights on the published STRmix™ results and the diagnostics used in PG software tools

## Footnotes

1 e.g. GeneMapper™

2 Our method displays high precision also for a simpler ‘Sample 2’ with log_10_ LR = 29.0144 *±* 0.00254.

## Notes

### Competing Interest Statement

GenoProof Mixture - a probabilistic genotyping software mentioned in the submitted manuscript - is developed by qualitype GmbH. Holger Schoenborn is one of the authors of this software.
Mateusz Susik is employed by Biotype GmbH. Biotype GmbH and qualitype GmbH are members of the Molecular Diagnostics Group consortium.

### Summary of Updates

Improved wording and preparation for a journal submission.

