## Supplementary Materials for "Hamiltonian Monte Carlo with strict convergence criteria reduces run-to-run variability in forensic DNA mixture deconvolution"

—

March 23, 2022

This Supplement contains:

- further explanation of the dropout model (Section 1),
- more detailed analysis of two mixtures of interest (Section 2),
- advise on how to implement the described HMC algorithm (Section 3),
- the list of stutter models used (Section 4),
- details about the experimental results (Section 5).

#### 1 Dropout model

Existing dropout models assume $P(\text{dropout}|x^l_{ar}) = P(z^l_{ar} < h^l_r | x^l_{ar})$, where $z^l_{ar}$ is the height of the raw peak before applying the analytical threshold [2, 5, 4]. Our log-normal model then states: $P(\text{dropout}|x^l_{ar}) = P\left(Z \leq \ln \frac{h^l_r}{x^l_{ar}}\right)$. For simplicity of notation, but without loss of generality, we assume in the following that a single peak-height variance parameter $c$ is used.

In early works, the standard deviation was assumed to be inversely proportional to the square root of the expected peak heights [12]. Since then, the model used by STRMix™ has evolved. In the current version, to the best of our knowledge, the standard deviation is assumed to be inversely proportional to

$$\sqrt{\frac{1000}{x^l_{ar}} + x^l_{ar}}\,. \tag{1}$$

We believe that this was done because the issue with assuming $X_{\text{old}} = \ln\left(\frac{O_a^l}{x_{ar}^l}\right) \sim \mathcal{N}\left(0, \frac{c^2}{x_{ar}^l}\right)$ is that

$$\lim_{x_{ar}^l \to 0} F_{X_{\text{old}}}\left(\ln \frac{h_r^l}{x_{ar}^l}\right) = \frac{1}{2}\,. \tag{2}$$

The Probability Density Function (PDF) $f_{X_{\text{old}}}\left(\ln \frac{h_r^l}{x_{ar}^l}\right)$ is a non-monotonic function of $x_{ar}^l$. It is increasing on $[0, b]$ and decreasing on $[b, \infty]$, where $b \in (0, h_r^l)$. Intuitively, the probability of a dropout event should be close to 1 if the expected peak height is close to 0 and a reasonable analytical threshold is used. This contradicts Eq. 2.

However, assuming $X_{\text{STRMix}^{\text{TM}}} \sim \mathcal{N}\left(0, \frac{c^2}{\frac{1000}{x_{ar}^l} + x_{ar}^l}\right)$ has the issue that:

$$\lim_{x_{ar}^l \to 0} \frac{1000}{x_{ar}^l} + x_{ar}^l = \infty\,. \tag{3}$$

In our work, we avoid this issue by regularizing the variance denominator using a shift of 1:

$$X_{\text{our}} \sim \mathcal{N}\left(0, \frac{c^2}{\frac{1000}{x_{ar}^l + 1} + x_{ar}^l}\right)\,. \tag{4}$$

While this solves the issues, it is not the only possibility. Another way of ensuring that the value of the CDF has the desired behaviour for low expected peak heights would be:

$$X_{\text{thresh}} \sim \mathcal{N}\left(0, \frac{c^2}{\max(h_r^l, x_{ar}^l)}\right)\,. \tag{5}$$

These different standard deviation models are plotted in Supplementary Fig. 1. Both Eqs. 4 and 5 exhibit the desired behaviour:

- The value of the CDF is 1 at 0.
- The value of the CDF is close (equal for Eq. 5) to 1/2 at the analytical threshold. This is expected for a log-normal distribution with mean 0 for the ratio of the observed and expected peak heights.
- The functions are monotonically decreasing.

### 2 Analysis of interesting cases

We analyse and discuss in detail two particularly interesting “odd” cases that serve to provide deeper insight into how the different inference methods work.

Supplementary Figure 1: Comparison of dropout standard deviation models for log-normal distributions. The STRMix™ models from Eq. 3 is not shown; it is almost identical to the function used in our work.

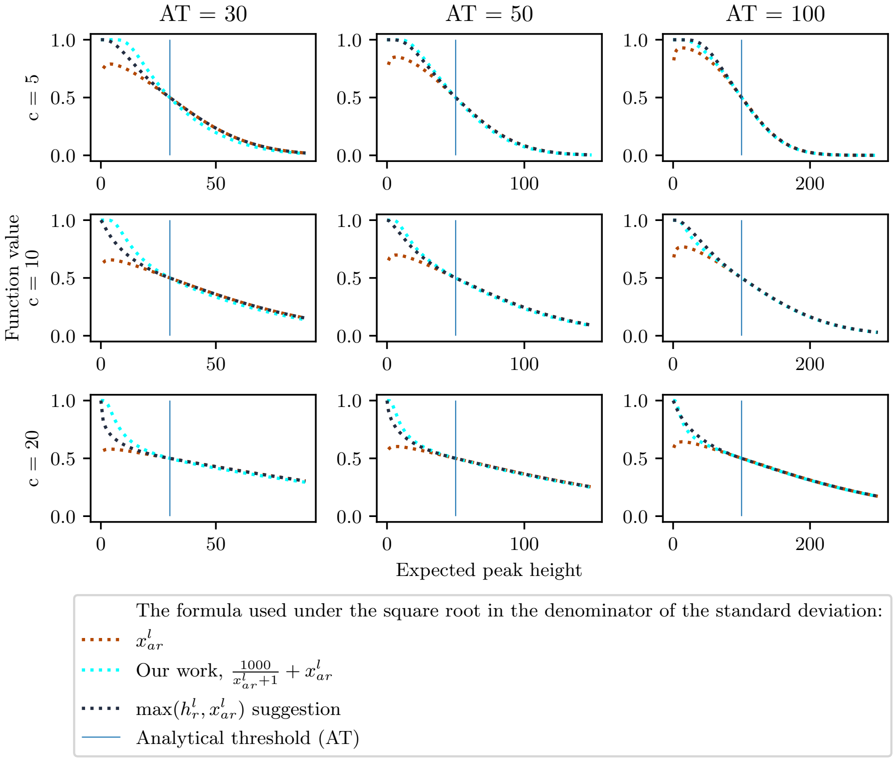


#### 2.1 LR larger than the inverse Random Match Probability (RMP): MIX13 Case 4

Without Balding-Nichols correction, LR must be smaller than 1/RMP [8]. In MIX13 Case 4, $\log_{10}(1/\mathrm{RMP}) = 20.187$ and our HMC inference method finds $\log_{10} \mathrm{LR} = 20.234$. We show that this apparent contradiction is indeed allowed when Balding-Nichols correction is used.

To see this, consider for example locus CSF1PO from this case. The known contributor's genotype is (10,12) and the suspect's genotype is (7,11). Then:

$$\frac{1}{\mathrm{RMP}} = \frac{1}{P(7,11|7,11 \text{ observed})} \tag{6}$$

$$= \frac{1.01 \cdot 1.02}{(0.01 + (0.99 \cdot 0.0025)) \cdot (0.01 + (0.99 \cdot 0.2995))} \approx 134.71\,. \tag{7}$$

MIX13 Case 4 is a two-contributor mixture. In the default scenario therefore is one unknown contributor in the defendant's hypothesis and one suspect in the prosecutor's hypothesis. Then, there is only one possible contributor order if the position of the known contributor is fixed. In such a case, the sub-source LR is equal to sub-sub-source LR. Consider a scenario in which the algorithm is able to correctly find the suspect's genotype with 100% confidence. This is the scenario that results in the highest possible LR for this locus. We denote this hypothetical maximum LR with Balding-Nichols correction at CSF1PO as $\mathrm{LR}_{\max}^{\mathrm{CSF1PO}}$. Then:

$$\mathrm{LR}_{\max}^{\mathrm{CSF1PO}} = \frac{1}{P(7,11|7,10,11,12 \text{ observed})} \tag{8}$$

$$= \frac{1.03 \cdot 1.04}{\mathrm{RMP} \cdot 1.01 \cdot 1.02} = \frac{1.0398}{\mathrm{RMP}} \approx 140.08\,. \tag{9}$$

Therefore, $\log_{10} \mathrm{LR}_{\max}^{\mathrm{CSF1PO}} > \log_{10}(1/\mathrm{RMP})$ is perfectly valid when Balding-Nichols correction is used. Following similar reasoning, we calculate the largest LR for the entire mixture as: $\log_{10} \mathrm{LR}_{\max} = 20.269$, and therefore correctly $\log_{10} \mathrm{LR}_{\max} > \log_{10} \mathrm{LR}$.

#### 2.2 Low precision: ProvedIt Sample 2b with contributor 30 as the suspect

We show that the low precision in this case is caused by an unlikely dropout event at locus D22S1045. The D22S1045 peak heights are given in Table 1. The suspect's genotype at this locus is (12,15). The suspect is the minor contributor, and the mixture has been prepared with a 4:4:1 ratio of material from the donors. Expert manual analysis might reveal that peaks 11, 15, and 17 come from the two major contributors. Peaks 10, 14, 16, and 18 might be explained by stutter. By looking only at the provided locus, the genotype of the minor contributor therefore remains unclear. Knowing the ratios of the donor materials, we see that all possibilities have to be considered, including all of the observed peaks and possible dropouts.

In Table 2, we provide evidence that the low precision in our results is entirely caused by this locus. We compare the two runs of our method that lead to the extremal results, namely the maximum observed $\log_{10} \mathrm{LR} = 13.061$ (named "max run") and the minimum 12.65 (named "min run").

Table 1: Peak heights at locus D22S1045 of ProvedIt Sample 2b. Allele 12 of the minor contributor has dropped out.

| **Allele** | 10 | 11 | 14 | 15 | 16 | 17 | 18 |
|---|---|---|---|---|---|---|---|
| **Peak height** | 57 | 2271 | 382 | 5195 | 428 | 2792 | 148 |

Excluding locus D22S1045, the extremal $log_{10}\mathrm{LR}$ are 20.581 and 20.578, restoring high precision. The very low LR at locus D22S1045 indicates that the suspect's genotype was included only among the genotype sets with low assigned probabilities. In Table 3 we present the most likely candidate genotype sets according to our algorithm that included the true minor contributor's

Table 2: A comparison of sub-sub-source per-locus likelihood ratios from the HMC runs that resulted in the largest and smallest LR, respectively. Likelihood ratios for ten loci of ProvedIt Sample 2 are presented. The observed precision on the whole mixture can be explained by the large deviation of the LRs at locus D22S1045. As discussed in the text, this locus provides evidence against the inclusion of the suspect, in contrary to most other loci.

| **Locus** | **LR max run** | **LR min run** | **ratio** |
|---|---|---|---|
| CSF1PO | 10.63 | 10.62 | ≈1 |
| D10S1248 | 8.18 | 8.2 | 0.998 |
| D12S391 | 97.42 | 97.42 | 1 |
| D13S317 | 0.46 | 0.46 | 1 |
| D16S539 | 2.58 | 2.58 | 1 |
| D18S51 | 20.19 | 20.27 | 0.996 |
| D19S433 | 612.3 | 612.3 | 1 |
| D1S1656 | 46.43 | 46.43 | 1 |
| D21S11 | 16.45 | 16.5 | 0.997 |
| **D22S1045** | **3.02e-08** | **1.18e-08** | **2.56** |

genotype, along with the corresponding genotype weights from both extremal runs. Maximum likelihood analysis of this case with our model reveals that the most likely expected peak height from the minor contributor for a dropped-out peak is 459.14. Given that locus D22S1045 is located on the red dye, an analytical threshold of 50 was used [9]. Under such circumstances, a dropout event is highly unlikely under our assumptions in the model. These results indicate that cases where the suspect's genotype is highly unlikely to appear in the deconvolution of some of the loci will result in decreased precision. We therefore also expect lower precision in cases where false contributors are assumed in the prosecutor's hypothesis.

We note, however, that simply ignoring D22S1045 is not a solution, because it introduces a bias into the analysis (even though this is sometimes done, for example see the results from participant L41.3A1 in the Supplementary Material of Ref. [3]). A possible solution would be to develop a new dropout model that is better suited for rare stochastic events. We leave research on this topic to future work.

Table 3: The three most likely genotype sets that explain the minor contributor's genotype: allele 15 and a drop-out at allele designation 12.

| | | | |
|---|---|---|---|
| **Major contributor 1** | (11,15) | (15,17) | (15,15) |
| **Major contributor 2** | (15,17) | (11,15) | (11,17) |
| **Minor contributor** | (15,Q) | (15,Q) | (15,Q) |
| **Weight max run** | 6.47e-10 | 1.23e-10 | 1.85e-10 |
| **Weight min run** | 2.1e-10 | 8.83e-11 | 4.64e-11 |

### 3 Implementation advice

We provide some useful hints and general advise for implementing our method in a software package. All of these are based on our own implementation experience and may apply to different platforms and software systems to various degrees.

#### 3.1 Constraining the parameter space

The continuous inference parameters have natural constraints:

- $c_s$ and $c_p$ have to be larger than 0;
- $t_r$ have to be larger that 0;
- $w_n$ have to sum up to 1 and all have to be non-negative;
- $d_{rn}$ have to be non-negative;
- $\alpha_r^l$ have to be larger than 0;

Hamiltonian Monte Carlo operates on the unconstrained domain $\mathbb{R}^N$. Therefore, we define a differentiable, invertible, element-wise transformation $b : \mathbb{R}^N \mapsto \mathbb{S}$ to the constrained parameter space $\mathbb{S}$ as:

- $b_c(x) = \mathrm{e}^x$ ;
- $b_{t_r}(x) = 30 + 1000 \ln(1 + \mathrm{e}^x)$ — we suggest to shift by a small constant (30 in our case) in order to avoid unstable gradients when the estimate of $t_r$ is very small. Scaling by 1000 normalises the magnitude of this parameter to be similar to that of the other parameters;
- $b_{w_n}(\mathbf{x}) = \frac{\exp([c(\mathbf{W}),0])}{\exp([c(\mathbf{W}),0])^\top J_{|\mathbf{w}|,1}}$ where:

  $$c(\mathbf{w})_n = \begin{cases} 0.1 x_n & \text{if } n \in K \\ 0.1 \ln\left(1 + \exp\left(x_{u_1} + \sum_{i \in [2, n-|K|]} \exp(x_{u_i})\right)\right) & \text{otherwise}, \end{cases} \tag{10}$$

  where $c(\mathbf{w})_n$ is the $n$-th element of the vector $c(\mathbf{w})$, $K$ is the set of indices of known contributors, and $x_{u_i}$ is the $i$-th element of the vector $\mathbf{x}$ that does not correspond to a known contributor. We index from 1, known contributors are always assigned the first indices, and the unknown contributor's weights follow. The vector $c(\mathbf{w})_n$ orders the unknown contributors in the minor-to-major order with an exception of the smallest unknown contributor, whose weight is added within the softmax operation by appending a zero. This avoids issues of label switching during the inference process. $J_{|\mathbf{w}|,1}$ is a vector of ones with the same length as vector $\mathbf{w}$.
- $b_{d_{rn}}(x) = \mathrm{e}^x$ ;
- $b_i(x) = 0.01\mathrm{e}^x$ ;

- $b_{\alpha^l_r}(x) = \mathrm{e}^x$ ;

This element-wise transformation and the corresponding corrections for the change of variables are applied to the model before HMC inference.

##### 3.2 The log-sum-exp trick

Given that many samples during inference have very low probability, we operate on log-probabilities. Then:

$$\begin{aligned}\log P(M|V) &= \log\left(P(M)\sum_j P(V|M,S_j)\right)\\ &= \log P(M) + \log\sum_j P(V|M,S_j)\,. \quad (11)\end{aligned}$$

In our implementation, the logarithm of the sum is omitted with the log-sum-exp-trick (LSE):

$$\log P(M) + \log\sum_j P(V|M,S_j) = \log P(M) + \mathrm{LSE}_j(\log P(V|M,S_j))\,, \quad (12)$$

where:

$$\mathrm{LSE}_j(\mathbf{x}) = \log\sum_j \exp(x_j)\,. \quad (13)$$

##### 3.3 Runtime optimization

We apply the following implementation optimisations for better overall runtime:

- use the Accelerated Linear Algebra (XLA) compiler of TensorFlow;
- initialise the HMC chains with reasonable parameter guesses. For this, we first run a maximum-likelihood estimation on a simplified surrogate model that only considers the sums of the peak heights at each locus to guess initial values for $\alpha^l_r$, $d_{rn}$, and $t'_r$. These initial values are then randomly perturbed (by parameter-specific scaled intensities) to provide different starting points for the different HMC runs;
- focus on limiting the number of divergent transitions while setting the hyper-parameters of the method [1]. Our measures include an adaptive schema for the initialisation of step-sizes of HMC and clipping the model gradients.

#### 4 Stutter models

For some loci, it has been shown that allele designation is not a good predictor of stutter ratio, and that instead Longest Uninterrupted Stretch (LUS) should be used [12]. For example, for locus TH01 it has been shown that the stutter ratio of allele designation 9.3 is very similar to that of allele designation 6 [12]. We therefore define:

$$\mathrm{LUS}_{\mathrm{TH01}}(n) = \begin{cases} n & \text{if } n \in \mathrm{N}, \\ n - 3.3 & \text{otherwise} . \end{cases} \tag{14}$$

For all other loci, we calculated LUS from the published data on variant alleles [11, 6, 7, 10]. The results are given in Table 4. Before modelling the ratios with linear models we translated the allele designations to numbers of expected repeats. For example, allele score 9.3 becomes 9.75 as there are 3 out of 4 nucleotides in the incomplete repeat.

| **kit** | **stutter type** | **locus** | **model type** | **intercept** | **coefficient** |
|---|---|---|---|---|---|
| AmpFℓSTR COfiler® | backward | D3S1358 | allele | -0.043 | 0.0063 |
| AmpFℓSTR COfiler® | backward | D16S539 | allele | -0.029 | 0.0066 |
| AmpFℓSTR COfiler® | backward | TH01 | LUS | -0.015 | 0.004 |
| AmpFℓSTR COfiler® | backward | TPOX | allele | -0.015 | 0.0033 |
| AmpFℓSTR COfiler® | backward | CSF1PO | allele | -0.043 | 0.0074 |
| AmpFℓSTR COfiler® | backward | D7S820 | allele | -0.042 | 0.0074 |
| AmpFℓSTR™SGM Plus™ | backward | D3S1358 | allele | -0.043 | 0.0063 |
| AmpFℓSTR™SGM Plus™ | backward | vWA | allele | -0.025 | 0.005 |
| AmpFℓSTR™SGM Plus™ | backward | D16S539 | allele | -0.029 | 0.0066 |
| AmpFℓSTR™SGM Plus™ | backward | D2S1338 | allele | -0.068 | 0.0065 |
| AmpFℓSTR™SGM Plus™ | backward | D8S1179 | allele | -0.029 | 0.0056 |
| AmpFℓSTR™SGM Plus™ | backward | D21S11 | allele | -0.024 | 0.0026 |
| AmpFℓSTR™SGM Plus™ | backward | D18S51 | allele | -0.032 | 0.0062 |
| AmpFℓSTR™SGM Plus™ | backward | D19S433 | allele | -0.081 | 0.0106 |
| AmpFℓSTR™SGM Plus™ | backward | TH01 | LUS | -0.015 | 0.004 |
| AmpFℓSTR™SGM Plus™ | backward | FGA | allele | -0.152 | 0.008 |
| AmpFℓSTR®Profiler Plus® | backward | D3S1358 | allele | -0.043 | 0.0063 |
| AmpFℓSTR®Profiler Plus® | backward | vWA | allele | -0.025 | 0.005 |
| AmpFℓSTR®Profiler Plus® | backward | D8S1179 | allele | -0.029 | 0.0056 |
| AmpFℓSTR®Profiler Plus® | backward | D21S11 | allele | -0.024 | 0.0026 |
| AmpFℓSTR®Profiler Plus® | backward | D18S51 | allele | -0.032 | 0.0062 |
| AmpFℓSTR®Profiler Plus® | backward | FGA | allele | -0.152 | 0.008 |
| AmpFℓSTR®Profiler Plus® | backward | D7S820 | allele | -0.042 | 0.0074 |
| AmpFℓSTR®Profiler Plus® | backward | D5S818 | allele | -0.002 | 0.0006 |
| AmpFℓSTR®Profiler Plus® | backward | D13S317 | allele | -0.004 | 0.0007 |
| GlobalFiler™ | backward | D10S1248 | allele | -0.052 | 0.0089 |
| GlobalFiler™ | backward | D16S539 | allele | -0.0465 | 0.009 |
| GlobalFiler™ | backward | CSF1PO | allele | -0.041 | 0.0086 |
| GlobalFiler™ | backward | D2S441 | allele | 0.0436 | 0 |
| GlobalFiler™ | backward | D19S433 | allele | -0.0552 | 0.0084 |
| GlobalFiler™ | backward | TH01 | LUS | -0.0221 | 0.006 |
| GlobalFiler™ | backward | D22S1045 | allele | -0.1094 | 0.0124 |
| GlobalFiler™ | backward | D7S820 | allele | -0.0434 | 0.0086 |
| GlobalFiler™ | backward | SE33 | allele | 0.0461 | 0.0018 |
| GlobalFiler™ | backward | D12S391 | allele | -0.0873 | 0.0086 |
| GlobalFiler™ | backward | D2S1338 | allele | -0.0192 | 0.0045 |
| GlobalFiler™ | backward | vWA | allele | -0.0917 | 0.0094 |
| GlobalFiler™ | backward | TPOX | allele | -0.0255 | 0.0054 |
| GlobalFiler™ | backward | D18S51 | allele | -0.0436 | 0.0074 |
| GlobalFiler™ | backward | D13S317 | allele | -0.0461 | 0.0082 |
| GlobalFiler™ | backward | D1S1656 | LUS | -0.0541 | 0.0099 |
| GlobalFiler™ | backward | D5S818 | allele | -0.0379 | 0.0083 |
| GlobalFiler™ | backward | D3S1358 | LUS | -0.0044 | 0.0069 |
| GlobalFiler™ | backward | D21S11 | LUS | 0.0027 | 0.006 |
| GlobalFiler™ | backward | FGA | allele | -0.0046 | 0.0032 |
| GlobalFiler™ | backward | D8S1179 | allele | 0.0251 | 0.0025 |
| GlobalFiler™ | forward | D22S1045 | allele | -0.0599 | 0.0062 |
| GlobalFiler™ | forward | D2S1338 | allele | -0.0028 | 0.0005 |
| GlobalFiler™ | forward | D16S539 | allele | -0.0039 | 0.001 |
| GlobalFiler™ | forward | D1S1656 | LUS | 0.0009 | 0.0007 |
| GlobalFiler™ | forward | D2S441 | allele | -0.0017 | 0.0008 |
| GlobalFiler™ | forward | D13S317 | allele | -0.0121 | 0.0017 |
| GlobalFiler™ | forward | D21S11 | allele | -0.0098 | 0.0007 |
| GlobalFiler™ | forward | SE33 | allele | 0.0085 | 0 |
| GlobalFiler™ | forward | CSF1PO | allele | -0.0047 | 0.001 |
| GlobalFiler™ | forward | D18S51 | allele | -0.0015 | 0.0007 |
| GlobalFiler™ | forward | D5S818 | allele | -0.0086 | 0.0016 |
| GlobalFiler™ | forward | D7S820 | allele | 0.0015 | 0.0005 |
| GlobalFiler™ | forward | D8S1179 | LUS | -0.0125 | 0.0019 |
| GlobalFiler™ | forward | D19S433 | LUS | 0.0065 | 0.0001 |

| | | | | | |
|---|---|---|---|---|---|
| GlobalFiler™ | forward | FGA | allele | -0.0045 | 0.0006 |
| GlobalFiler™ | forward | D12S391 | allele | 0.008 | 0 |
| GlobalFiler™ | forward | D10S1248 | allele | -0.0015 | 0.0006 |
| GlobalFiler™ | forward | vWA | allele | 0.0032 | 0.0003 |
| GlobalFiler™ | double backward | D22S1045 | allele | -0.0077 | 0.0011 |
| GlobalFiler™ | double backward | D12S391 | allele | -0.0094 | 0.0009 |
| GlobalFiler™ | double backward | SE33 | allele | 0.0035 | 0.0002 |
| GlobalFiler™ | double backward | D2S1338 | allele | 0.0008 | 0.0003 |
| GlobalFiler™ | double backward | D18S51 | allele | -0.0045 | 0.0008 |
| GlobalFiler™ | double backward | D21S11 | LUS | -0.0052 | 0.0011 |
| GlobalFiler™ | double backward | D1S1656 | LUS | -0.0036 | 0.0008 |
| GlobalFiler™ | double backward | D8S1179 | LUS | -0.0078 | 0.0013 |
| GlobalFiler™ | double backward | D10S1248 | allele | -0.0036 | 0.0008 |
| AmpFℓSTR™Identifiler™Plus | backward | D8S1179 | allele | 0.0202 | 0.0031 |
| AmpFℓSTR™Identifiler™Plus | backward | D7S820 | allele | -0.0439 | 0.0087 |
| AmpFℓSTR™Identifiler™Plus | backward | TH01 | LUS | -0.0245 | 0.0064 |
| AmpFℓSTR™Identifiler™Plus | backward | D13S317 | allele | -0.0475 | 0.0086 |
| AmpFℓSTR™Identifiler™Plus | backward | D16S539 | allele | -0.0514 | 0.0096 |
| AmpFℓSTR™Identifiler™Plus | backward | TPOX | allele | -0.0252 | 0.0057 |
| AmpFℓSTR™Identifiler™Plus | backward | CSF1PO | allele | -0.0356 | 0.008 |
| AmpFℓSTR™Identifiler™Plus | backward | D3S1358 | LUS | -0.0089 | 0.0076 |
| AmpFℓSTR™Identifiler™Plus | backward | D2S1338 | allele | -0.0167 | 0.0046 |
| AmpFℓSTR™Identifiler™Plus | backward | D19S433 | LUS | -0.0308 | 0.0085 |
| AmpFℓSTR™Identifiler™Plus | backward | D5S818 | allele | -0.0297 | 0.0077 |
| AmpFℓSTR™Identifiler™Plus | backward | FGA | allele | -0.0123 | 0.0036 |
| AmpFℓSTR™Identifiler™Plus | backward | vWA | allele | -0.0969 | 0.01 |
| AmpFℓSTR™Identifiler™Plus | backward | D21S11 | LUS | -0.0023 | 0.0066 |
| AmpFℓSTR™Identifiler™Plus | backward | D18S51 | allele | -0.0432 | 0.0079 |
| AmpFℓSTR™Identifiler™Plus | forward | D8S1179 | LUS | -0.0004 | 0.0007 |
| AmpFℓSTR™Identifiler™Plus | forward | D13S317 | allele | 0.0007 | 0.0005 |
| AmpFℓSTR™Identifiler™Plus | forward | D3S1358 | allele | 0.0065 | 0 |
| AmpFℓSTR™Identifiler™Plus | forward | D16S539 | allele | -0.0016 | 0.0007 |
| AmpFℓSTR™Identifiler™Plus | forward | CSF1PO | allele | -0.0069 | 0.0012 |
| AmpFℓSTR™Identifiler™Plus | forward | FGA | allele | -0.0006 | 0.0005 |
| AmpFℓSTR™Identifiler™Plus | forward | TH01 | LUS | 0.0044 | 0.0001 |
| AmpFℓSTR™Identifiler™Plus | forward | TPOX | allele | 0.0059 | 0 |
| AmpFℓSTR™Identifiler™Plus | forward | D21S11 | LUS | -0.0044 | 0.0011 |
| AmpFℓSTR™Identifiler™Plus | forward | D5S818 | allele | -0.0079 | 0.0015 |
| AmpFℓSTR™Identifiler™Plus | forward | D7S820 | allele | -0.0009 | 0.0006 |
| AmpFℓSTR™Identifiler™Plus | forward | D2S1338 | allele | 0.0097 | 0 |
| AmpFℓSTR™Identifiler™Plus | forward | D18S51 | allele | -0.0009 | 0.0007 |
| AmpFℓSTR™Identifiler™Plus | forward | vWA | allele | 0.0101 | 0 |
| AmpFℓSTR™Identifiler™Plus | forward | D19S433 | LUS | 0.0045 | 0.0003 |
| AmpFℓSTR™Identifiler™Plus | double backward | D8S1179 | allele | 0.0048 | 0 |
| AmpFℓSTR™Identifiler™Plus | double backward | D3S1358 | LUS | 0.0064 | 0.0001 |
| AmpFℓSTR™Identifiler™Plus | double backward | D7S820 | allele | 0.0003 | 0.0004 |
| AmpFℓSTR™Identifiler™Plus | double backward | D18S51 | allele | -0.0031 | 0.0007 |
| AmpFℓSTR™Identifiler™Plus | double backward | D13S317 | allele | -0.0036 | 0.0007 |
| AmpFℓSTR™Identifiler™Plus | double backward | D19S433 | LUS | 0.0056 | 0.0003 |
| AmpFℓSTR™Identifiler™Plus | double backward | FGA | allele | 0.0009 | 0.0004 |
| PowerPlex®16 | backward | TH01 | LUS | -0.0443 | 0.0099 |
| PowerPlex®16 | backward | D18S51 | allele | -0.0484 | 0.0086 |
| PowerPlex®16 | backward | Penta E | allele | -0.0195 | 0.0043 |
| PowerPlex®16 | backward | D5S818 | allele | -0.0444 | 0.0103 |
| PowerPlex®16 | backward | D7S820 | allele | -0.0447 | 0.0102 |
| PowerPlex®16 | backward | D16S539 | allele | -0.0517 | 0.0115 |
| PowerPlex®16 | backward | CSF1PO | allele | -0.0524 | 0.0109 |
| PowerPlex®16 | backward | vWA | allele | -0.2013 | 0.0174 |
| PowerPlex®16 | backward | FGA | allele | -0.0852 | 0.0078 |
| PowerPlex®16 | backward | D3S1358 | LUS | -0.0523 | 0.0135 |
| PowerPlex®16 | backward | Penta D | allele | -0.0201 | 0.0034 |
| PowerPlex®16 | backward | D8S1179 | LUS | -0.0166 | 0.0072 |
| PowerPlex®16 | backward | TPOX | allele | -0.0287 | 0.0065 |
| PowerPlex®16 | backward | D21S11 | LUS | -0.0348 | 0.0118 |
| PowerPlex®16 | backward | D13S317 | allele | -0.0518 | 0.0105 |
| PowerPlex®16 | forward | D3S1358 | LUS | -0.0012 | 0.0015 |
| PowerPlex®16 | forward | D21S11 | allele | -0.0282 | 0.0015 |
| PowerPlex®16 | forward | D18S51 | allele | -0.0055 | 0.001 |
| PowerPlex®16 | forward | Penta E | allele | 0.0023 | 0.0003 |
| PowerPlex®16 | forward | D5S818 | allele | 0.0013 | 0.0012 |
| PowerPlex®16 | forward | D13S317 | allele | -0.0157 | 0.003 |
| PowerPlex®16 | forward | D7S820 | allele | -0.0018 | 0.0016 |
| PowerPlex®16 | forward | CSF1PO | allele | -0.0174 | 0.0028 |
| PowerPlex®16 | forward | vWA | allele | -0.0179 | 0.0016 |
| PowerPlex®16 | forward | FGA | allele | -0.0161 | 0.0013 |
| PowerPlex®16 | forward | D16S539 | allele | -0.0121 | 0.0024 |
| PowerPlex®16 | forward | D8S1179 | LUS | 0.0013 | 0.0008 |
| PowerPlex®16 | double backward | D18S51 | allele | -0.0063 | 0.0008 |
| PowerPlex®16 | double backward | Penta E | allele | -0.0007 | 0.0004 |
| PowerPlex®16 | double backward | CSF1PO | allele | -0.0035 | 0.0008 |
| PowerPlex®16 | double backward | FGA | allele | 0.0218 | 0.0002 |
| PowerPlex®16 | double backward | D3S1358 | LUS | -0.0053 | 0.0013 |
| PowerPlex®16 | double backward | D8S1179 | LUS | -0.0106 | 0.0014 |
| PowerPlex®16 | double backward | D16S539 | allele | -0.011 | 0.0017 |

| | | | | | |
|---|---|---|---|---|---|
| PowerPlex®16 | double backward | D7S820 | allele | -0.0112 | 0.0018 |
| PowerPlex®16 | double backward | D5S818 | allele | 0.0085 | 0.0001 |

Table 4: Stutter models used in the benchmarks based on either allele designation or Longest Uninterrupted Stretch (LUS) with parameters as indicated.

### 5 Benchmark results

We complement the results from the main text with additional details and explanations, and we investigate possible sources of variability from different software versions.

#### 5.1 Precision study

Given the large number of benchmarks we performed, we provide the means and standard deviations in Table 6. Detailed raw results are available upon request.

| Analysis method | Metric | Mean | StdDev |
|---|---|---|---|
| A | Max GR of 1.05 | 4.5281 | 0.01296 |
| A | Mean GR of 1.2 | 4.5388 | 0.0216 |
| A | STRMix™-like of 1.05 | 4.5366 | 0.04115 |
| B | Max GR of 1.05 | 6.2564 | 0.0164 |
| B | Mean GR of 1.2 | 6.2753 | 0.0328 |
| B | STRMix™-like of 1.05 | 6.3005 | 0.05179 |
| C | Max GR of 1.05 | 4.5364 | 0.01319 |
| C | Mean GR of 1.2 | 4.5429 | 0.02675 |
| C | STRMix™-like of 1.05 | 4.546 | 0.03649 |
| D | Max GR of 1.05 | 5.7633 | 0.01652 |
| D | Mean GR of 1.2 | 5.7608 | 0.03352 |
| D | STRMix™-like of 1.05 | 5.8161 | 0.04472 |

Table 5: Means and standard deviations of the results presented in Fig. 2 of the main text.

The previously published results had been obtained using different versions of STRMix™ ranging from version 2.4.05 to version 2.5.11. One could argue for a possible source of variability due to the differences in the source code between these software versions. In Supplementary Figure 2, we therefore compare the results obtained with different versions of STRMix™ for ProvedIt Sample 1, Analysis Method D. We choose this case because it has the largest sample size. We use a two-sided Kolmogorov-Smirnov test to test for the null hypothesis that all results come from the same underlying distribution. The number of samples obtained with the different software versions are: 79 results for 2.4.08, 31 for 2.5.11, 9 for 2.4.05, and 7 for 2.4.06. Comparing versions 2.4.08 and 2.5.11, the p-value of the test is 0.27, making it impossible to reject the null hypothesis of identical underlying distributions. We therefore conclude that the variability caused by different software versions of STRmix™ is not statistically significant.

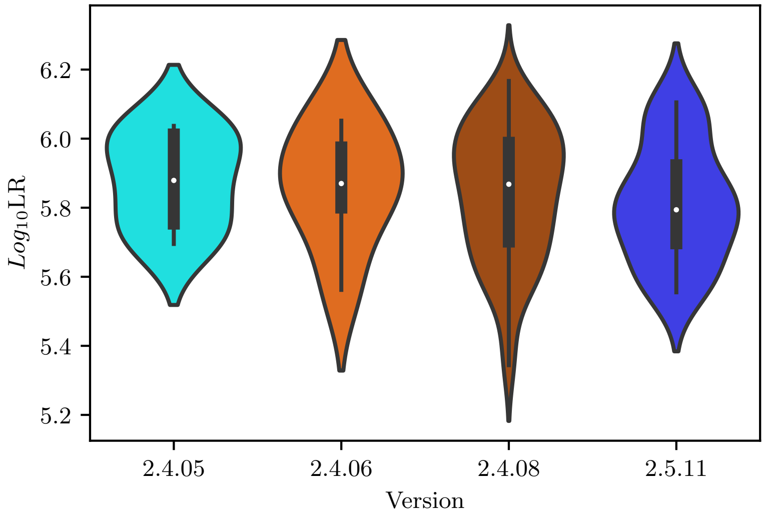


Supplementary Figure 2: Violin plots of $\log_{10}$ LR obtained with different software versions (given on the x-axis) of STRMix™ on ProvedIt Sample 1, Analysis Method D. The null hypothesis of all results coming from the same distribution cannot be rejected (two-sided Kolmogorov-Smirnov test, p-value 0.27, N=79 and 31).

#### 5.2 Computationally challenging mixtures

| Sample | Suspect | | | | | | | | | | |
|---|---|---|---|---|---|---|---|---|---|---|---|
| 1b | 44 | 1.426 | 1.418 | 1.383 | 1.458 | 1.402 | 1.405 | 1.452 | 1.438 | 1.414 | 1.463 |
| 1b | C58A˙Asian | -2.093 | -1.957 | -1.821 | -2.127 | -1.868 | -2.043 | -2.194 | -2.088 | -1.962 | -2.2 |
| 1b | 46 | 17.261 | 17.241 | 17.264 | 17.281 | 17.249 | 17.358 | 17.306 | 17.28 | 17.261 | 17.288 |
| 1b | C75H˙Hisp | -1.303 | -1.21 | -1.127 | -1.323 | -1.153 | -1.278 | -1.378 | -1.316 | -1.216 | -1.366 |
| 1b | 47 | 5.139 | 5.137 | 5.155 | 5.135 | 5.147 | 5.185 | 5.148 | 5.14 | 5.148 | 5.123 |
| 1b | 45 | 3.169 | 3.129 | 3.071 | 3.181 | 3.097 | 3.11 | 3.197 | 3.175 | 3.135 | 3.205 |
| 1b | ZT79899˙Hisp | -2.754 | -2.65 | -2.529 | -2.806 | -2.578 | -2.714 | -2.856 | -2.757 | -2.652 | -2.867 |
| 1b | C61B˙AA | -2.164 | -2.057 | -1.912 | -2.226 | -1.973 | -2.087 | -2.296 | -2.186 | -2.049 | -2.311 |
| 2b | 30 | 12.697 | 12.757 | 12.999 | 12.81 | 12.811 | 12.839 | 12.85 | 13.005 | 12.898 | 12.7 |
| 2b | 31 | 17.876 | 17.871 | 17.884 | 17.864 | 17.873 | 17.885 | 17.88 | 17.89 | 17.885 | 17.877 |
| 2b | 32 | 17.873 | 17.868 | 17.88 | 17.861 | 17.869 | 17.881 | 17.877 | 17.887 | 17.881 | 17.874 |
| 3 | 33 | 2.791 | 2.748 | 2.767 | 2.692 | 2.824 | 2.75 | 2.718 | 2.713 | 2.763 | 2.725 |
| 3 | 30 | 8.612 | 8.593 | 8.638 | 8.626 | 8.625 | 8.576 | 8.603 | 8.622 | 8.606 | 8.599 |
| 3 | 32 | 10.556 | 10.433 | 10.461 | 10.472 | 10.557 | 10.448 | 10.495 | 10.504 | 10.478 | 10.479 |
| 3 | 34 | 8.69 | 8.7 | 8.703 | 8.675 | 8.676 | 8.702 | 8.697 | 8.693 | 8.719 | 8.686 |
| 3 | C42C˙Cauc | -2.117 | -2 | -2.186 | -1.908 | -2.172 | -2.099 | -1.939 | -1.98 | -2.133 | -1.934 |
| 3 | WT51359˙Cauc | -6.448 | -6.083 | -6.752 | -5.827 | -6.567 | -6.255 | -6.09 | -6.251 | -6.488 | -6.123 |
| 3 | GT38089˙Cauc | -2.816 | -2.697 | -2.917 | -2.584 | -2.885 | -2.754 | -2.664 | -2.732 | -2.806 | -2.684 |
| 3 | JT51484˙AA | -2.695 | -2.49 | -2.881 | -2.312 | -2.746 | -2.611 | -2.413 | -2.554 | -2.705 | -2.472 |
| 4 | C45A˙Asian | -1.698 | -1.683 | -1.678 | -1.666 | -1.721 | -1.691 | -1.683 | -1.698 | -1.672 | -1.688 |
| 4 | 48 | 0.14 | 0.164 | 0.165 | 0.157 | 0.159 | 0.157 | 0.175 | 0.156 | 0.161 | 0.157 |
| 4 | PT84541˙Hisp | -0.629 | -0.687 | -0.629 | -0.632 | -0.589 | -0.622 | -0.616 | -0.655 | -0.636 | -0.661 |
| 4 | GT37166˙AA | -0.293 | -0.328 | -0.294 | -0.295 | -0.289 | -0.305 | -0.3 | -0.312 | -0.287 | -0.3 |
| 4 | 49 | 1.95 | 1.957 | 1.925 | 1.939 | 1.923 | 1.934 | 1.919 | 1.949 | 1.944 | 1.933 |
| 4 | 50 | 7.873 | 7.846 | 7.916 | 7.927 | 7.902 | 7.896 | 7.903 | 7.882 | 7.88 | 7.881 |
| 4 | C94C˙Cauc | -1.799 | -1.838 | -1.774 | -1.767 | -1.782 | -1.79 | -1.779 | -1.822 | -1.776 | -1.826 |
| 4 | 30 | 2.734 | 2.739 | 2.71 | 2.708 | 2.722 | 2.727 | 2.731 | 2.73 | 2.713 | 2.718 |

Table 6: Raw $\log_{10}$ LR results of our benchmarks for the computationally challenging mixtures. Cases that resulted in LR=0 for all 10 runs are omitted.

### References


[1] Michael Betancourt. Diagnosing suboptimal cotangent disintegrations in hamiltonian monte carlo, 2016.

[2] Øyvind Bleka, Geir Storvik, and Peter Gill. EuroForMix: An open source software based on a continuous model to evaluate STR DNA profiles from a mixture of contributors with artefacts. *Forensic Science International: Genetics*, 21:35–44, March 2016.

[3] Jo-Anne Bright, Kevin Cheng, Zane Kerr, Catherine McGovern, Hannah Kelly, Tamyra R. Moretti, Michael A. Smith, Frederick R. Bieber, Bruce Budowle, Michael D. Coble, Rashed Alghafri, Paul Stafford Allen, Amy Barber, Vickie Beamer, Christina Buettner, Melanie Russell, Christian Gehrig, Tacha Hicks, Jessica Charak, Kate Cheong-Wing, Anne Ciecko, Christie T. Davis, Michael Donley, Natalie Pedersen, Bill Gartside, Dominic Granger, MaryMargaret Greer-Ritzheimer, Erick Reisinger, Jarrah Kennedy, Erin Grammer, Marla Kaplan, David Hansen, Hans J. Larsen, Alanna Laureano, Christina Li, Eugene Lien, Emilia Lindberg, Ciara Kelly, Ben Mallinder, Simon Malsom, Alyse Yacovone-Margetts, Andrew McWhorter, Sapana M. Prajapati, Tamar Powell, Gary Shutler, Kate Stevenson, April R. Stonehouse, Lindsey Smith, Julie Murakami, Eric Halsing, Darren Wright, Leigh Clark, Duncan A. Taylor, and John Buckleton. STRmix™ collaborative exercise on DNA mixture interpretation. *Forensic Science International: Genetics*, 40:1–8, May 2019.

[4] John Buckleton. The probability of dropout and drop-in.

[5] Frank M. Götz, Holger Schönborn, Viktoria Borsdorf, Anne-Marie Pflugbeil, and Dirk Labudde. GenoProof mixture 3—new software and process to resolve complex DNA mixtures. *Forensic Science International: Genetics*, 6:e549–e551, December 2017.

[6] Marielle Heinrich, Heike Felske-Zech, Bernd Brinkmann, and Carsten Hohoff. Characterisation of variant alleles in the STR systems d2s1338, d3s1358 and d19s433. *International Journal of Legal Medicine*, 119(5):310–313, June 2005.

[7] Maria Victoria Lareu, Maria del Carmen Pestoni, Francisco Barros, Antonio Salas, and Angel Carracedo. Sequence variation of a hypervariable short tandem repeat at the d12s391 locus. *Gene*, 182(1-2):151–153, December 1996.

[8] Mark W. Perlin. Genotype likelihood ratio distributions and random match probability : Generalization , calculation and application. 2017.

[9] Sarah Riman, Hari Iyer, and Peter M. Vallone. Examining performance and likelihood ratios for two likelihood ratio systems using the PROVEDIt dataset. *PLOS ONE*, 16(9):e0256714, September 2021.

[10] B. Rolf, M. Schürenkamp, A. Junge, and B. Brinkmann. Sequence polymorphism at the tetranucleotide repeat of the human beta-actin related pseudogene h-beta-ac-psi-2 (ACTBP2) locus. *International Journal of Legal Medicine*, 110(2):69–72, February 1997.

[11] C. M. Ruitberg. STRBase: a short tandem repeat DNA database for the human identity testing community. *Nucleic Acids Research*, 29(1):320–322, January 2001.

[12] Duncan Taylor, Jo-Anne Bright, and John Buckleton. The interpretation of single source and mixed DNA profiles. *Forensic Science International: Genetics*, 7(5):516–528, September 2013.
